# Patterns of gene-body-methylation conservation and its divergent association with gene expression in Pigeonpea and Soybean

**DOI:** 10.1101/2020.03.11.987339

**Authors:** Alim Junaid, N.K. Singh, Kishor Gaikwad

**Author notes:** Corresponding author: Dr. Kishor Gaikwad.

## Abstract

Pigeonpea and Soybean are closely related agronomically important legumes that diverged ∼23 million years ago followed by a whole genome duplication event in Soybean. However, the impact of divergence on their epigenomes is not known. Comparative epigenomics is a powerful tool to understand shaping of epigenomes and its functional consequences during evolution. In this study, we investigated the conservation and divergence of Gene body methylation (GbM) and gene-expression patterns between the two species. We compared their genome-wide DNA methylation landscape at single-base resolution and studied the influence of GbM on duplicate gene retention. Our results indicate that during the divergence of Pigeonpea and Soybean, the effects of methylation on gene expression evolved in a heterogeneous manner. GbM features-slow evolution rate and increased length remained conserved in Pigeonpea and Soybean. We also found that contrary to high CG-methylation conservation, non-CG methylated genes remained diverged during evolution due to transposons insertion in adjacent sequences. The methylation differences in paralogs were associated with expression divergence. Notably, we found that variably methylated marks are less likely to target evolutionary conserved sequences. Finally, our findings identify conservation of nitrogen-related genes during evolution, which are promising candidates for epi-alleles for crop improvement.

## Introduction

DNA methylation is an evolutionary conserved epigenetic mechanism in plants and animals that involves methylation of cytosine bases at position 5 of the cytosine ring and regulates gene expression and genome stability (Robertson, 2005; Slotkin & Martienssen, 2007). In contrast to animals, DNA methylation in plants occurs in all cytosine contexts i.e., CG, CHG and CHH (where H denotes A, T or C).

Different features of the genome are subjected to DNA methylation, based on its function. Regulation of gene expression involves occurrence of DNA methylation at different positions within a gene-promoter or the transcribed gene body. Methylation in promoter regions is profoundly associated with transcriptional repression (Domcke et al., 2015; X. Zhang et al., 2006; Zhu, Wang, & Qian, 2016) but they have been shown to activate transcription in certain instances(Lang et al., 2017; Liu et al., 2015). Gene body methylation (GbM) occurs primarily in mCG context within the coding regions of genes and is excluded from the transcriptional start and termination sites(Takuno & Gaut, 2013). Although biological functions of GbM have not been fully discovered, it has been proposed to play a role in enhancing the efficiency of pre-mRNA splicing and regulate gene homeostasis by preventing transcription from cryptic promoters (Regulski et al., 2013; Takuno & Gaut, 2012; Zilberman, Coleman-Derr, Ballinger, & Henikoff, 2008). Notably, GbM has been proposed to have evolutionary consequences. A comparative analysis of GbM between Rice and *Brachypodium*, that diverged at least 40-53 million years ago, revealed a conservation of GbM between orthologs that was further proposed to have a functional implication of targeting a subset of genes for methylation(Takuno & Gaut, 2013). A similar trend was also shown to occur between *Arabidopsis lyrata* and *Arabidopsis thaliana* that diverged around 26 million years ago(Takuno & Gaut, 2013). GbM has been shown to target long and slowly evolving house-keeping genes, and this feature is conserved in angiosperms with an exception in Poaceae family where gene body methylated genes have been shown to evolve at a faster rate (Niederhuth et al., 2016; Takuno & Gaut, 2012, 2013; X. Zhang et al., 2006). Furthermore, GbM is also proposed to impact evolutionary properties of genes in grasses and other monocots, such as their nucleotide landscape particularly bimodal GC3 distribution(Carels & Bernardi, 2000; Serres-Giardi, Belkhir, David, & Glemin, 2012; Takuno & Gaut, 2013).

Evolutionary mechanisms such as genome duplication followed by divergence are fundamental to the evolution of angiosperm. Since the genomes of angiosperms are prone to polyploidization arising from whole genome duplication events (WGD), they have also evolved a tendency for diploidization (to prevent predictive extinction) and for duplicated gene loss over long evolutionary period (Coghlan, Eichler, Oliver, Paterson, & Stein, 2005; Otto & Whitton, 2000; Paterson, Freeling, Tang, & Wang, 2010; Wolfe, 2001). Duplicated genes may face two evolutionary fates- (1) biased retention of both copies owing to neo-functionalization by accumulating neutral mutations or gene balance via fractionation to single copy (Lewis 1951, Ohno 1970) (2) Selective loss of one of the paralogous genes. However, the knowledge on the influence of the epigenome on these evolutionary events is very limited. Recent advances in genome and epigenome sequencing have shed light on WGD events in different plants and their epigenetic landscape. It has been found that *Glycine max* got subjected to a recent WGD event with no immediate diploidization. Consequently around three quarter of genes of comprise of duplicated genes which are enriched in CG gene body methylation and show relatively higher expression levels compared to singleton genes(K. D. Kim et al., 2015; K. D. Kim, Shin, Van, Kim, & Lee, 2009; Schmutz et al., 2010). Another study in Cassava also reported positive correlation between gene body methylation and expression levels in the duplicated gene pairs (H. Wang et al., 2015). Although there are developing evidences of the influence of methylation on evolutionary fates of duplicated genes, comparative analysis of DNA methylation between closely related species, particularly crops will more exemplify its impact of duplicated gene retention or loss and establish a link between selection of preferential agronomic traits during evolution. Recent studies have revealed conservation of gene body methylation between orthologs of potato and tomato (L. Wang et al., 2018) and between Soybean and Common bean (K. D. Kim et al., 2015), but more comprehensive knowledge is required towards this end.

Pigeonpea (2n=22) and Soybean are agronomically important legume crops belonging to millettioid subgroup of the Fabaceae family. These two crops diverged around 20-30 million years ago(Lavin, Herendeen, & Wojciechowski, 2005) followed by the occurrence of more recent genome duplication event in Soybean (13 million years ago) (Schmutz et al., 2010). Despite these events, Pigeonpea and Soybean share high levels of synteny, notably at every chromosome (Varshney et al., 2011). Therefore, Pigeonpea and Soybean offer an attractive system to investigate the evolutionary functions of DNA methylation during and after polyploidization and speciation.

In this study we provide insights into methylome dynamics occurring during evolution and speciation by comparing the DNA methylation landscapes of Pigeonpea and Soybean at single base-pair resolution. We specifically investigate roles and the balance of and CG Gene body methylation (GbM) conservation and divergence during evolution by exclusive analysis of CG GbM patterns in both species. We further compare GbM within orthologous and paralogous gene between Pigeonpea and Soybean to determine the dynamics of DNA methylation patterns after speciation and Soybean specific polyploidy.

## Methods

### Plant materials and Growth conditions

Pigeonpea lines 11A (male sterile), 303R (fertile) and 11A X 303R (hybrid) were used for experiments and analysis(Junaid et al., 2018). Plants were grown in green house under natural conditions at the Indian Agricultural Research Institute New Delhi, India. Genotypes were verified by pollen fertility test using 2% acetocarmine dye as described previously (Junaid et al., 2018).

### DNA isolation and BS-Seq library preparation

Immature young buds were harvested from 3-month-old plants. Two biological replicates were used for each of the three lines. High molecular weight DNA was extracted using CTAB method with slight modification. The pair end BS-seq libraries were prepared using illumina true seq kit according to manufacturer’s instructions. Briefly, 1-5 ug of DNA was sheared to 100-300 bp using Covaris-M220 and ensued DNA fragments were end repaired by adding dA at 3′-end prior to the ligation of true seq methylated adapters. The adapter ligated DNA fragments were treated with bisulfite using EZ DNA Methylation-Gold kit (D5005) and sequenced using illumina Hi-seq 2000 platform following manufacture’s instruction.

### BS-Seq data analysis

Raw reads containing adapters and low-quality bases at the 5’ and 3’ end was cleaned using Cutadapt and Trim galore (https://github.com/FelixKrueger/TrimGalore). High quality reads were mapped to the reference genome of Pigeonpea(Varshney et al., 2011) and Soybean (Schmutz et al., 2010) using bsmap at default parameter (Xi & Li, 2009). The methylation status of each mapped cytosine bases was extracted using *methratio*.*py* and weightage methylation level was calculated. The metaplots analysis of genes and transposons were performed by dividing 2kb flanking and gene body region into equal size bin and then methylation level was calculated for each bin and plotted in ggplot2. Previously published data (SRR5850382, SRR1628892) of Soybean methylome was used for analysis.

### Gene body methylation determination

The GbM of each gene were determined according to (Takuno & Gaut, 2012) with a modified strategy. Firstly, genes were classified into T.E and Non-T.E category based on transposons insertion/absence and then CG, CHG and CHH methylation level in CDS region was calculated for both gene categories separately. We used a minimum coverage cut-off of 3X for CG and CHG and 10X for CHH context to calculate body methylation for genes. Afterwards binomial probability distribution was applied to calculate one-tailed P value based on divergence of CG methylation level of gene from average genomic methylation level.

### Identification of differentially methylated region (DMRs)

To identify DMRs between three genotypes, Methyl kit (Akalin et al., 2012) pipeline was used. In brief, genome was tiled into 100 bp windows and weightage methylation level was calculated for each window and pair-wise test was performed. The filtering criteria was also adopted to exclude ambiguous cytosine resulting from incomplete conversion or sequencing error. We used cytosine with minimum 10X coverage and at least three cytosine for CG and CHG whereas six cytosines for CHH to define 100bp window. We used logistic regression model to calculate P-values which was further adjusted to FDR using SLIM method for final DMR identification. The windows with methylation difference with at least 20% and q-value < 0.01 for all the three contexts were considered as candidate DMRs.

### RNA-seq library preparation

Total RNA from the bud of pigeonpea hybrid line was extracted using Sigma (E4913) according to the manufacturer’s instructions. The RNA-seq libraries were prepared in two biological replicates using the standard TrueSeq RNA sample preparation kit (Illumina) and sequenced in 100bp pair end on illumina Hi Seq 2000 platform. For soybean transcriptome, we used publicly available data of NJCMS1A line (SRR1752076).

### RNA-Seq data analysis

Raw reads were filtered for adapter removal, the high-quality reads were aligned to Soybean and Pigeonpea reference genomes using tophat2 (D. Kim et al., 2013) at default parameters. The ensued aligned reads were provided for transcript assembly and quantification of transcript abundance using cufflinks. The transcript expression was determined in terms of fragments per kb of exon per million fragments mapped (FPKM). Transcripts were annotated against Non-redundant NCBI data base using Blast2GO (https://www.blast2go.com). The GO enrichment analysis was performed using AgriGO (http://bioinfo.cau.edu.cn/agriGO/) and enriched terms were defined with Fisher’s exact test and False discovery rate (FDR) correction by Bonferroni method (Du, Zhou, Ling, Zhang, & Su, 2010).

#### Transposons identification

We used a combinational approach to identify robust set of transposons in Pigeonpea and Soybean. The soybean transposons library was downloaded from soya base. While, repeat modeler was used to construct a denovo repeat library of pigeonpea genome. Afterwards, Repeat Masker was run at default parameters using repeat libraries to identify transposons. Transposons less than 100 b.p were discarded from subsequent analysis. In addition to this, LTR digest and LTR harvest were also used to identify full length LTRs. The LTR data sets generated from both approaches were further compared and overlapped LTRs were considered for further analysis.

### Identification of homologous genes

Duplicated and syntenic gene pairs between pigeonpea and soybean were identified by using MCScanX(Y. Wang et al., 2012). The proteome sequences of both species were downloaded from NCBI and reciprocal blastp approach was adopted with parameter of e-value 1e-10 and output was generated in -m 8 format. The resulting tab delimited output was provided to MCScanX and subsequently collinear regions were identified. The genes were classified into different duplicated categories using command *duplicate_gene_classifier*. Protein sequences of homologous genes were aligned and resulting alignment output was converted into axt format using clustaw2 (Thompson, Higgins, & Gibson, 1994) and paraAT (Zhang et al., 2012). Then, Ka and Ks values were calculated by employing Nei & Gojobori method (Nei & Gojobori, 1986) using Ka-Ks calculator(Z. Zhang et al., 2006).

## Results

### Comparative landscape of DNA methylation reveals higher genome-wide methylation levels in Pigeonpea relative to Soybean

We characterized and compared the methylomes of the floral buds of Pigeonpea lines 11A (male sterile) and 303R (fertile) and their F1 hybrid using two biological replicates, at single base pair resolution using whole genome bisulfite sequencing. In total 330 million reads were generated for all the three lines. After trimming the adaptors and low-quality reads, 72-84% cleaned reads were mapped to the Pigeonpea reference genome (Varshney et al., 2011) (Table S1). The bisulfite conversion efficiency was calculated by aligning reads to the unmethylated chloroplast and lambda genome. Thereafter, methylation levels in mCG, mCHG and mCHH contexts between biological replicates were compared in 2kb windows to check the variability within six sequenced libraries. A minimum coverage for cytosines in symmetric sites were considered to be at least four, whereas for asymmetric site it was increased to ten. We found high correlation in symmetric sites (r=0.94-0.9, CG; r= 0.92-0.94, CHG) and relatively lower correlation in asymmetric sites (r= 0.71-0.86) (Supporting Information Fig. S1). We also observed a bimodal distribution of methylation at symmetric sites (Supporting Information Fig. S1). However, methylation levels in majority of CHH sites were found to be highly reduced. A comparison of global DNA methylation patterns between the three genotypes of Pigeonpea revealed overall high mCG and mCHG weightage methylation levels in 303R line relative to 11A and F1 but higher mCHH methylation in 11A followed by F1 and 303R (Fig. 1A).

**Figure 1.**
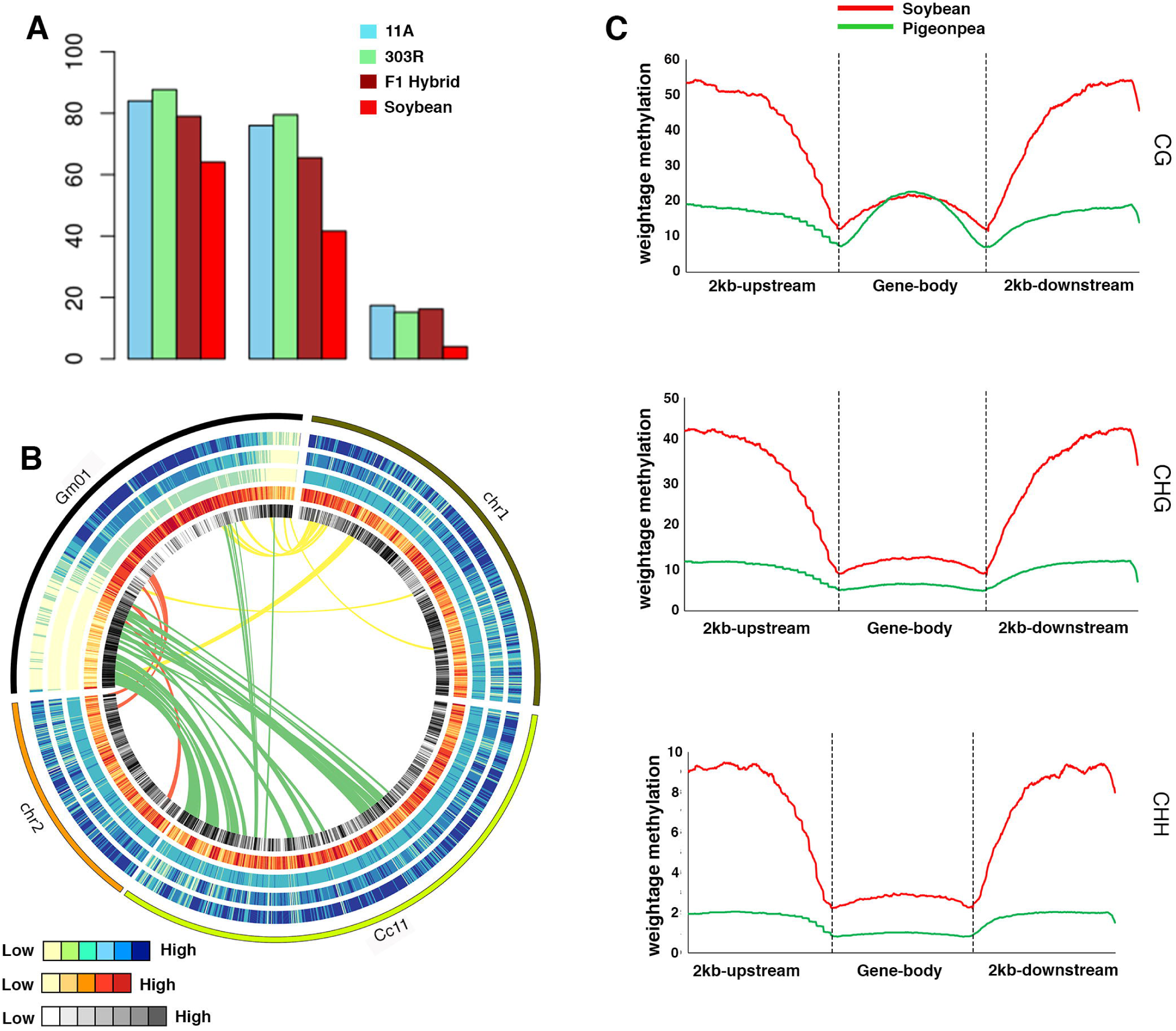
Comparative analysis of whole genome DNA methylation patterns of Pigeonpea and Soybean. (A) Comparison of genome-wide DNA weightage methylation levels of flower buds in three different genotypes (11A, 303R and F1 hybrid) of pigeonpea and Soybean. (B) A circos plot showing the DNA methylation landscape and syntenic association of homologous gene pairs in Soybean (Gm01) and Pigeonpea (Chr1, Chr2, Chr11). Heatmap shows CG, CHG and CHH methylation levels, transposons density and gene density (outer to inner circle). Red and black colors indicate high transposons and gene densities, respectively. Blue color indicates high methylation levels and yellow color indicates low methylation level. (C) Meta-plots showing mCG, mCHG and mCHH methylation pattern in gene body and flanking regulatory regions in Pigeonpea and Soybean (+2kb upstream from TSS and - 2kb downstream from TTS sites). Cc: *Cajanus cajan* (Pigeonpea); Gm: *Glycine max* (Soybean). The numbers indicate the respective chromosome numbers.

In-order to gain insights into the global methylome dynamics occurring during speciation and evolution, we compared genome-wide average methylation patterns between Pigeonpea and its closely related species Soybean, that diverged ∼20-30 million years ago (Varshney et al., 2011). We found much higher levels of weightage methylation in Pigeonpea than Soybean in all the mC contexts (Fig 1A). Genome wide landscapes of mCG and mCHG showed a positive correlation with transposon-density (r= 0.23; CG, r= 0.24; CHG P-value < 2.2e-16) and a negative correlation with the gene-density (r= −0.56; CG, r= −0.58; CHG P-value < 2.2e-16) in Pigeonpea (Fig. 1B). Chromosome wide distribution analysis of CG, CHG and CHH methylation surprisingly revealed an unexpected pattern for CHH methylation in Pigeonpea. Transposons rich regions showed a weak although significant negative association (r= −0.07; P-value =1.488e-11) and gene-dense regions showed a reduced positive association (r= 0.02; P-value = 0.04) with CHH methylation levels. However, unlike Pigeonpea the transposons rich centromeric regions showed relatively high methylation levels than the gene-dense telomeric regions of the chromosomes, in all three mC contexts in Soybean.

Lastly, we compared the CG, CHG and CHH weightage methylation levels of protein coding genes and their regulatory regions (2kb upstream and downstream) in Pigeonpea and Soybean. Genes intersecting with the transposons in gene body region were removed from subsequent metaplot analysis. The metaplots showed maximum distribution of CG, CHG and CHH methylation levels in the gene flanking regions, followed by a continuous drop with minima around TSS and TTS sites (Fig. 1C). Methylation levels increased continuously when moving away from these sites. We also observed a CG methylation peak within the gene body regions of both the species although it was relatively more prominent in Soybean than Pigeonpea. In contrast, CHG and CHH metaplots were marked by absence of this peak and methylation levels were uniform throughout the gene body. Unsurprisingly, consistent with whole genome methylation levels, Pigeonpea exhibited a higher methylation level in all three mC contexts in flanking as well as gene body regions relative to Soybean (Fig 1C).

### Comparative methylation landscape of Pigeonpea and Soybean TE regions

Plant genomes are enriched with transposable elements (TE elements) that generally exist in an epigenetically silenced state regulated via RdDM pathway(Fultz, Choudury, & Slotkin, 2015; Nuthikattu et al., 2013). To understand the evolutionary dynamics of TE methylation, we performed a detailed comparative analysis of CG, CHG and CHH methylation levels in different types of TEs in Pigeonpea and Soybean. All the TE types displayed highest methylation levels in CG context in both Pigeonpea and Soybean. We found overlapping TE body methylation levels in CG context for DNA elements and Retrotransposons (Copia, Gypsy) in Pigeonpea and Soybean (Fig. 2A, D, J) suggesting a robust maintenance of CG methylation levels in TE body during evolution. However, TE body methylation levels in CHG context were slightly higher in Pigeonpea than the Soybean (Fig. 2 B, E H, K). Furthermore, in both CG and CHG contexts, methylation levels in upstream and downstream regulatory sequences of DNA elements, GYPSY and Copia were comparable in Pigeonpea and Soybean. LINES transposons displayed a relatively higher CG and CHG methylation in the flanking regulatory sequences in Pigeonpea (Fig 2 G, H). Strikingly, for all TEs, CHH methylation throughout the TE body, upstream and downstream regulatory sequences was found to be significantly higher (4X) in Pigeonpea relative to Soybean suggesting that during evolutionary divergence of Soybean from Pigeonpea, evolutionary forces dampened the role of CHH methylation in regulating TE activity (Fig. 2C, F, I, L). Further investigation of the transposons in Pigeonpea genomes revealed the enrichment of short length transposons which have either low or high CG and CHG methylation level whereas long transposons were consistently remained in high methylated state. In contrast to CG and CHG, CHH methylation increases with transposons length (Supporting Information Fig. S2).

**Figure 2.**
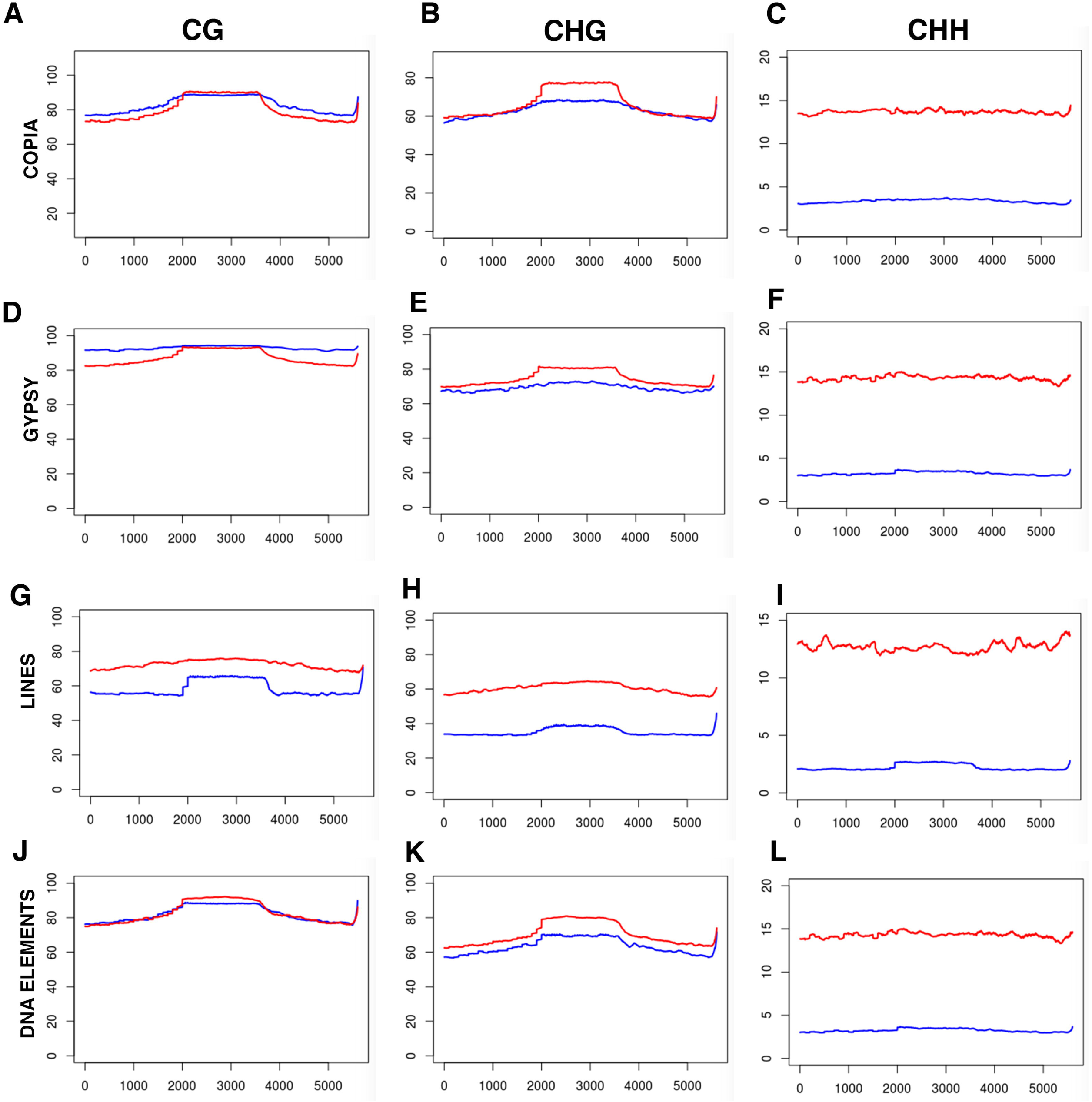
Comparative profile of DNA methylation variation across transposon regions in Pigeonpea and Soybean. (A-D) Metaplots showing the distribution of mCG, mCHG and mCHH along the two different family of transposons, class I (DNA elements) and class II transposons (Copia, Gypsy and Lines) in Pigeonpea (green line) and Soybean (red line). The flanking regulatory sequences (2kb upstream and 2 kb downstream) and gene body regions were divided into equal sized 20 bins and weightage methylation level were calculated for each bin.

### Extensive analysis of gene body methylation in Pigeonpea and Soybean

In-order to execute an intricate comparative analysis of genic methylation patterns between Pigeonpea and Soybean, genes were categorized based on the insertion or absence of transposons (TE-genes and non-TE genes respectively here after). We found a much lower proportion of TE-genes (17.3% and 15.8%) than non-TE genes (82.6% and 84.1%) in both Pigeonpea and Soybean (Fig. 3A). Thereafter, weightage methylation levels were calculated for both gene categories. In line with the predictions that transposons insertion contributes to increased methylation levels, we found high levels of methylation within the gene body of TE-genes relative to non-T.E genes in all mC contexts in both species (Supporting Information Fig. S3).

**Figure 3.**
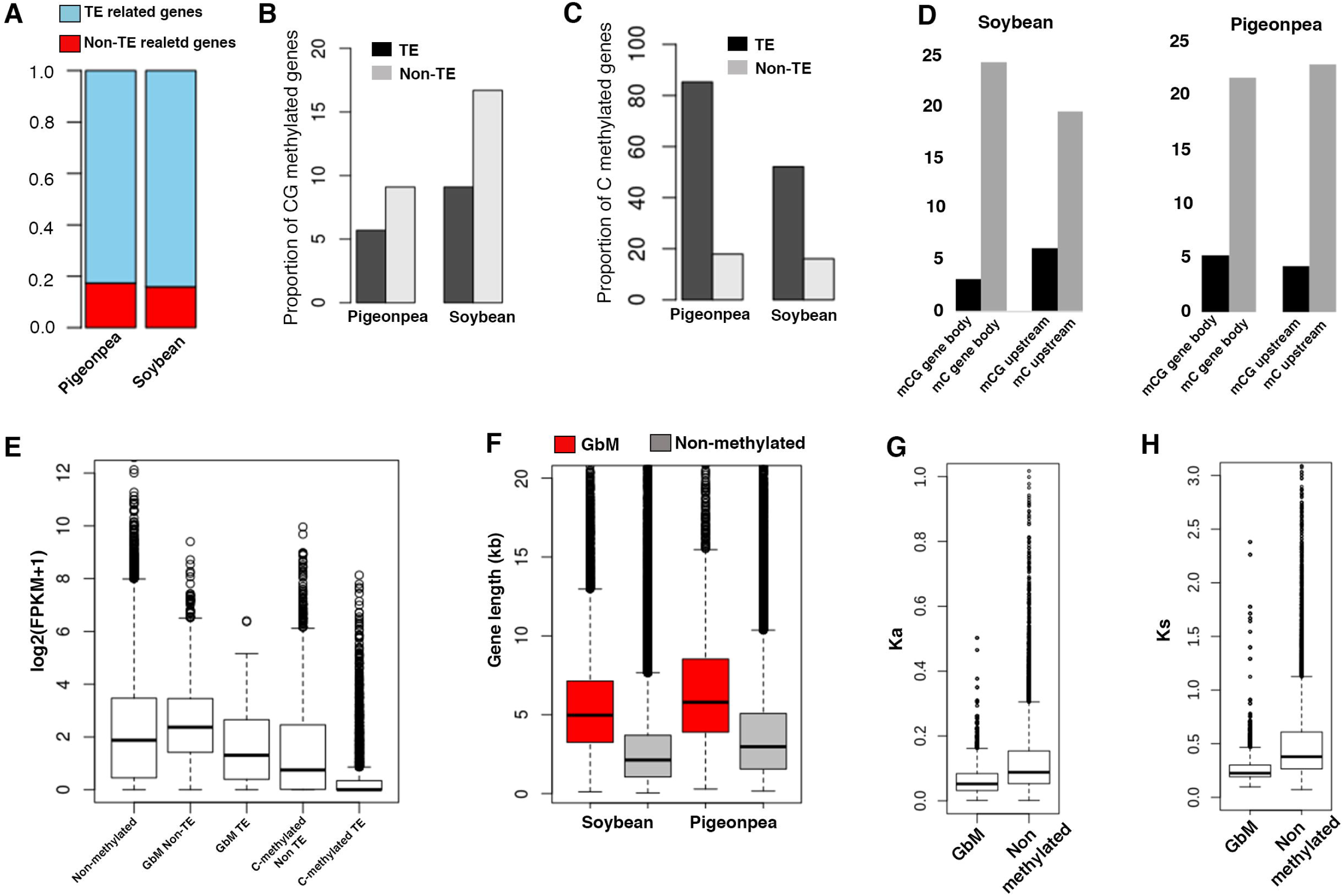
Characterization of distinct features of CG body and non-CG methylated genes. A. Classification of Pigeonpea and Soybean genes based on transposon insertion within gene body region. B. Fraction of (a) mCG (b) non-CG body methylated in the context of transposons insertion. The significance in distribution of CG body methylated genes was tested using Chi square test; p-value < 2.2e-16. C. Box plots showing longer average length of CG body methylated (red) genes than the un-methylated (grey) in both species. D. Percentage of transposons coverage in 2kb upstream and body region showing high abundance of transposons in non-CG methylated genes in both species. E. Boxplots showing normalized expression (log2(FPKM+1)) values of each gene category in Pigeonpea. The significance difference in expression level was calculated using Wilcoxon rank sum test: (P-values details in main text) F. Box plots showing gene length in GbM genes and non-methylated genes in Pigeonpea and Soybean. Note higher length of GbM genes in both species relative to non-methylated genes. G. Ka distribution of mCG body methylated and unmethylated ortholog pairs. CG body methylated genes showed low divergence relative to un-methylated genes. H. Ks distribution of mCG body methylated and unmethylated ortholog pairs. CG body methylated genes showed low divergence relative to un-methylated genes.

Next, in order to investigate the potential role of CG body methylation (GbM) and their conservation during evolution, TE-genes and non-TE genes were separately classified into CG-body (GbM) and C-methylated genes based on their genic background methylation level using probabilistic approach (Takuno & Gaut, 2012). Genes which were significantly enriched in CHG and CHH methylation with PCHG and PCHH <0.05 were categorized as C-methylated genes. GbM genes were identified with PCG <0.05 and methylated exclusively in CG context. We identified a total of 2608 (8.5%), 2343 (8.8%), 2763 (10%) and 8466 (15.5%) body methylated genes in 11A, 303R, hybrid and in Soybean respectively. The distribution of GbM genes in TE and non-T.E genes was found to be significantly different in both species (P-value < 2.2e-16; Chi square test). Among GbM genes, significant proportion in Pigeonpea (2245, 9.7% of total) and Soybean (7673, 16.7% of total) were non-T.E-genes, while only a smaller fraction of T.E genes (5.7% in Pigeonpea and 9.1% in soybean of total) were body methylated in both species (Fig.3B).

### GbM gene features are conserved between Pigeonpea and Soybean

Earlier studies in *Arabidopsis* and a few species of Poaceae family have demonstrated a positive correlation between gene lengths or number of exons and CG body methylation, where significantly body methylated genes were longer than the lowly methylated genes (Seymour & Gaut, 2019; Takuno & Gaut, 2012). Also, since GbM genes are more prone to spontaneous mutations due to frequent deamination of C to T (Bird, 1980; Pfeifer, 2006), it is expected that such genes would evolve at a faster rate to consolidate for potential deleterious effects of such mutations. Notably several studies have found that GbM genes evolve slowly (Takuno & Gaut, 2012, 2013; Takuno, Seymour, & Gaut, 2017). However, how the GbM features evolved during the divergence of Pigeonpea from Soybean are not known. Therefore, we investigated these specific features in Pigeonpea and Soybean GbM genes. We found that average length of GbM genes was 7.3 and 5.6 kb in Pigeonpea and Soybean (Fig. 3F) respectively, which was interestingly found to be about two-fold higher than the lowly methylated gene in both species. To decipher the evolution rate among GbM genes in Pigeonpea and Soybean, Synonymous (Ks) and Non-synonymous (Ka) substitution rates were calculated. We found that Ka (0.069; CG methylated, 0.12; Non methylated) and Ks (0.30; CG methylated, 0.49; Non-methylated) were significantly low in CG body methylated genes (P-value < 2.2e-16, Wilcoxon rank sum test) than the non-methylated genes (Fig. 3G, H).

### Association of TE enrichment and C-methylated genes

We observed that T.E-gene comprehended a high number of C-methylated genes where-in 56-84% of the genes were significantly methylated at multiple contexts in Pigeonpea and Soybean (Fig. 3C). We also found a high coverage of TEs in upstream as well as gene body regions of C-methylated genes in both species (Fig 3D). Since TE insertion in genes is presumed to disrupt their expression (Hollister & Gaut, 2009), we closely investigated the relationship between gene expression and transposons insertion in CG and mC gene categories of Pigeonpea (Fig. 3E). We found that overall, GbM genes showed high expression compared to mC genes irrespective of transposons insertion in Pigeonpea. However, GbM genes with TE insertion showed slightly reduced expression than the non-TE GbM genes, suggesting that GbM might influence gene expression in a specific way under distinct conditions. Further investigations revealed that mC genes with transposons inserted in their body region showed a significantly (Wilcoxon rank sum test; P-value < 2.2e-16) reduced expression among different genes category. These results suggest that transposons insertion within the genic regions may affect gene expression by disseminating methylation signatures in the 5’ upstream regulatory elements and that this mechanism is more favorable/predominant in the mC genes.

### DNA methylation exerts heterogeneous effects on gene expression in Pigeonpea and Soybean

A conclusive association between CG body methylation and gene expression remains obscure (Cokus et al., 2008; Hollister et al., 2011; Lister et al., 2008; Yang, Takuno, Waters, & Gaut, 2011). In-order to gain extensive insights into the relationship between DNA methylation and global gene expression pattern from an evolutionary context, transcriptome of Pigeonpea and Soybean were characterized. Intriguingly, we found a negative correlation (ρ= −0.15, P-value < 2.2e-16) of gene expression and CG methylation levels in Pigeonpea (Fig 4A), contrary to a positive correlation (ρ = 0.27, P-value < 2.2e-16) of gene body methylation and expression in Soybean (Fig 4D). Moderately CG methylated genes (30-50%) tend to have highest expression among all categories, which suggest that intermediate body methylation may have a positive impact on gene expression (Fig. 4A). In contrast to CG methylation, CHG (Fig. 4B, E) and CHH (Fig. 4C, F) methylation showed a negative association with gene expression in both species. Overall, these results indicate that during the divergence of Pigeonpea and Soybean, the effects of methylation on gene expression evolved in a heterogeneous manner, that may be attributed to species specific evolution of gene regulatory mechanism.

**Figure 4.**
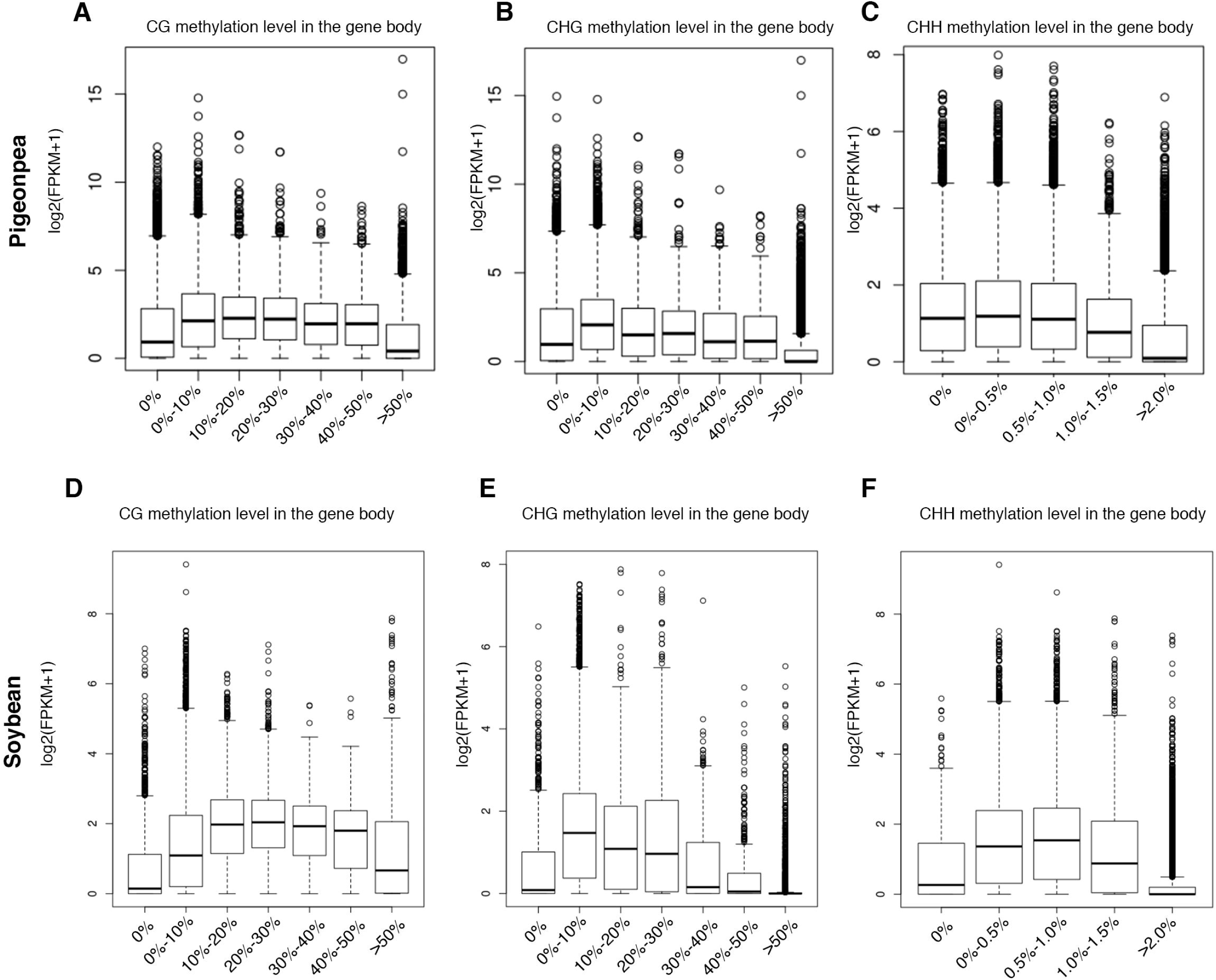
Correlation between genic methylation and expression levels. (A-F) Pigeonpea (A-C) and Soybean (D-F) genes classified based on mCG, mCHG and mCHH methylation levels displayed along x-axis and their normalized expression levels plotted on y-axis. Genes with CG and CHG methylation level 0% were classified as un-methylated genes and the remaining genes were divided into six groups. The 7^th^ group contained genes with >50% CG and CHG methylation in both species. Different criteria were applied to classify CHH methylated genes. Genes with CHH methylation >2% were kept in last (highly methylated) group. Note a negative correlation of CG methylation and expression in all three contexts in Pigeonpea. A positive correlation in mCG gene body methylation and expression was found whereas the methylation in mCHG and mCHH genes were negatively associated with expression.

### CG gene body methylation is conserved in Pigeonpea and Soybean homologous genes

In-order to determine the evolutionary changes in DNA methylation patterns post the events of speciation and Soybean*-*specific polyploidy, GbM within the orthologous and paralogous genes between Pigeonpea and Soybean was compared. We identified a total of 46,538 out of 95,711 genes (48.3%) as orthologous gene pairs between Soybean and Pigeonpea and 35,050 out of 54,324 (64%) and 4636 out of 29,192 (16%) paralogous gene pairs in Soybean and Pigeonpea respectively. For an explicit CG methylation comparative analysis, gene pairs containing C-methylated genes (*P*_CHG_ <0.05 or *P*_CHH_<0.05) were excluded, based on earlier report (K. D. Kim et al., 2015). We found a highly positive correlation (r=0.53, r=0.69 and r=0.41) in CG methylation levels across orthologous gene pairs as well as paralogous gene pairs in Soybean and Pigeonpea, respectively (Fig. 5 A, C, E). Notably, scatter plots revealed relatively high CG methylation levels in Pigeonpea genes, consistent with the whole genome methylation levels. In contrast, we observed a relatively low positive correlation in non-CG methylation levels between the orthologs (r= 0.1, CHG and r= 0.08, CHH) and paralogs (Pigeonpea: r=0.19 CHG and r=0.1; CHH, Soybean: r=0.3, CHG and r=0.21) (Fig. 5B, D, F, Supporting Information Fig. S4). A further meticulous analysis revealed that 4.3 % of genes pairs (1632 out of 37109) in collinear blocks showed significant CG body methylation in both species, while only small fraction, 0.9% of gene pairs showed significant C-methylation in both copies of genes. Above, we showed that a large proportion of genes are non-methylated in Pigeonpea and Soybean (Fig 3B). Consistent with our findings, a large proportion of orthologs, ∼69.9% were found to be non-methylated (20826 of 29757 in Soybean and 11742 of 16781 in Pigeonpea). These results reflect that during evolution, methylation state of the non-methylated genes remained robust. A very low percentage, ∼7.5% (2259 of 29757) of Soybean non-CG methylated genes were found orthologous in Pigeonpea and 7.4%-7.8% of Pigeonpea CG methylated genes (1320/16781, F1 hybrid; 1248/16781, 11A and 1268/16781, 103R) were found orthologous in Soybean. Next, to dissect methylation patterns at species level, we analyzed methylation status in synteny genes separately in Soybean and Pigeonpea. We found that ∼19.10% (5685 of 29757) of Soybean and 8.42%-10.31% of Pigeonpea CG methylated genes (1731/16781; F1 hybrid, 1413/16781; 11A and 1590/16781; 103R) were present within the colinear orthologous regions. All-together, our findings indicate CG gene body methylation patterns remained conserved between orthologs and paralogs of Soybean and Pigeonpea. Further, since our earlier analysis revealed that C-methylated genes have high TE coverage (Fig.3 D, E) our findings that non-CG body methylated genes are not highly conserved during evolution could possibly relate to a high frequency of TE insertion within C-methylated orthologs. Additionally, the observed methylation patterns also indicate that the evolutionary forces favored relatively high conservation of CG body methylation in Soybean than in Pigeonpea during speciation.

**Figure 5.**
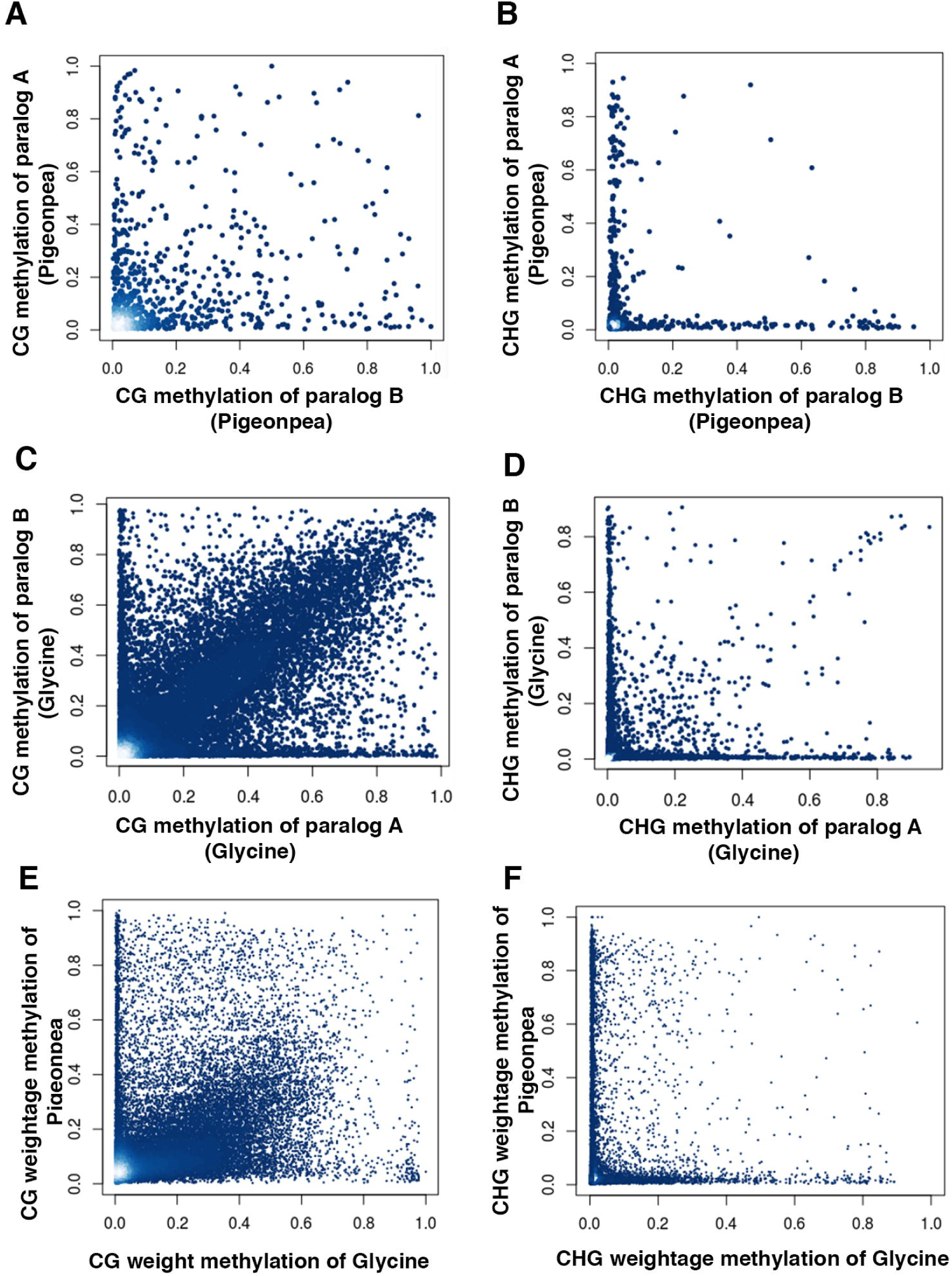
Correlation of DNA methylation level of homologous gene pairs in Pigeonpea and Soybean. (A-F) Pairwise comparison of CG and CHG weightage methylation levels in paralogous gene pairs (A-D) and orthologous gene pairs (E-F). Spearman correlation coefficient was calculated to show association (details in main text).

We next sought to determine the functional categories of GbM orthologous genes between Pigeonpea and Glycine using Gene Ontology (GO) enrichment analysis. The most significant GO terms included functional categories associated with nitrogen compound biosynthetic processes, nitrogen compound transport, nitrogen compound metabolic processes as well as gene categories belonging to carbohydrate derivatives transport (Table S2). Given the fundamental role of nitrogen (N) in plant growth, reproduction and protein content of seeds especially in legumes, it is compelling that GbM orthologs of the two most important legume crops that diverged ∼30 million years ago are enriched for gene categories belonging to nitrogen metabolism, transport and biosynthesis.

### Analysis of methylation patterns between paralogous genes reveals covariation in methylation and expression divergence

Plant genomes are repeatedly restructured during evolution, resulting in genes to acquire different origins. Therefore, we next aimed to trace the evolution of methylation patterns within the genes of different origins in Pigeonpea and Soybean. We found 64% of genes were segmentally duplicated (WGD) in Soybean, contrary to only a small proportion in Pigeonpea (16.35%). These observations comply with the recent genome duplication event in Soybean. The abundance of CG-methylated WGD genes (18.4%) was slightly higher than the singleton genes (11%) in Soybean. In contrast, the percentage of CG-methylated WGD genes was lower (9.3%) than the singleton genes (11%) in Pigeonpea (Fig. 6A). Since single gene duplicates (SGDs) were highly proportioned in majority of mC genes in both species (Fig. 6A) and previous studies have demonstrated the roles of TE-related mechanisms in the origin of SGDs (Drouin & Dover, 1990; Jiang, Bao, Zhang, Eddy, & Wessler, 2004) we examined the TE coverage within and SGD and WGD and found a high TE coverage in SGD than the WGD genes (Fig 6B, C).

**Figure 6.**
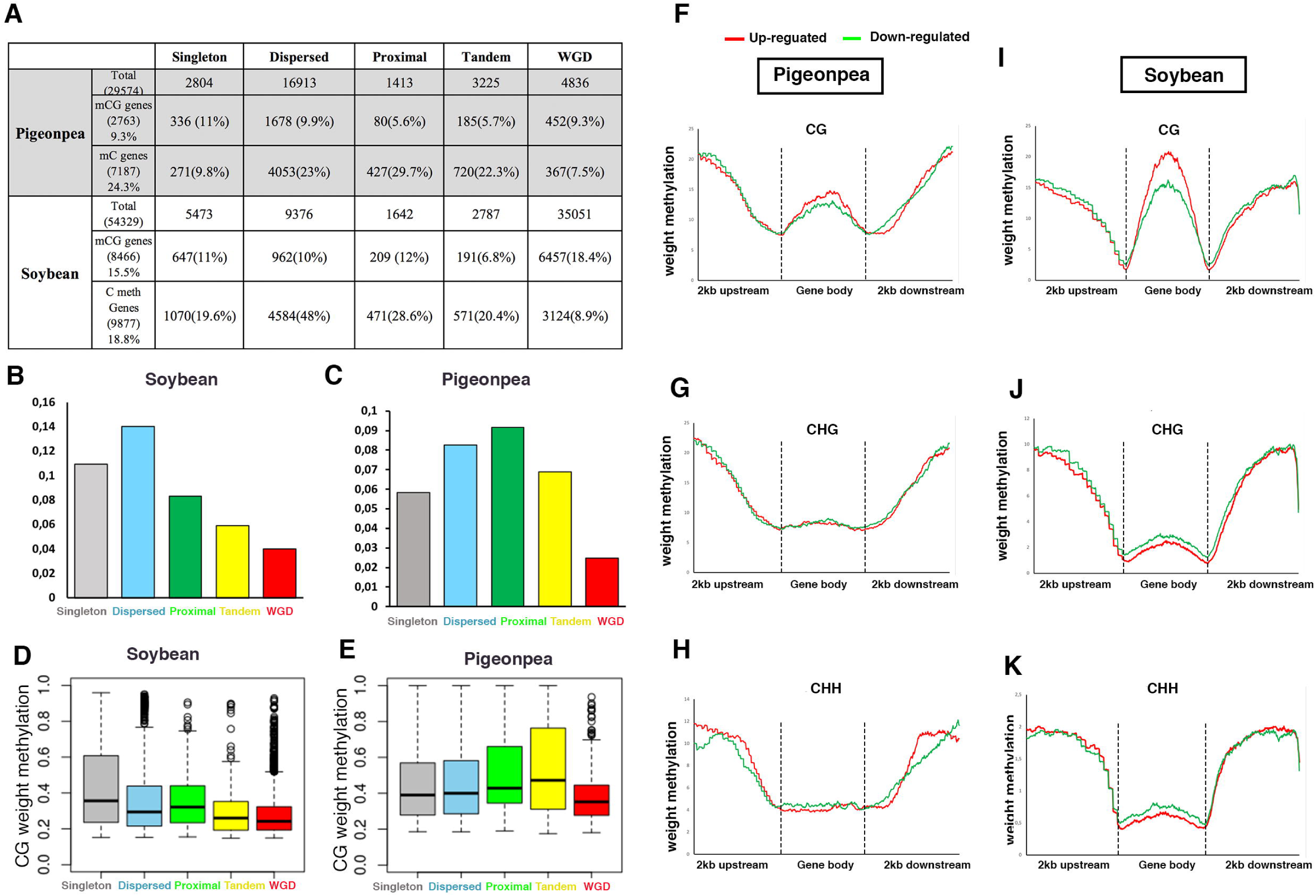
Gene body methylation and expression analysis in four different categories of whole genome duplicated (WGD) and single copy genes. A. Distribution of methylated genes in WGD genes of Pigeonpea and Soybean. B. Boxplots showing comparison of weightage methylation and expression pattern in WGD CG body methylated genes in Pigeonpea (A, C) and Soybean (B, D). (E) Meta-plots showing methylation patterns in gene-flanking (2 kb upstream and 2 kb downstream) and gene body regions of highly (log2FC >2) and lowly (log2FC < −2) expressed duplicate gene pairs. Red and green lines represent highly and lowly expressed genes respectively. The significance test was performed using Wilcoxon rank-sum test. (P-values in main text).

We next calculated and compared the CG weightage methylation levels of duplicated (SGD and WGD) and singleton GbM genes. In Pigeonpea, proximal mCG genes displayed the highest methylation levels. WGD mCG genes showed significantly low methylation levels relative to singleton genes (Wilcoxon rank sum test P-value < 0.008). Strikingly, WGD mCG genes methylation levels were also found to be significantly lower in comparison to other duplicated gene categories (proximal (P-value= 3.915e-05); tandem (P-value= 2.53e-05) and dispersed (P-value = 6.836e-10). We also observed a statistically significant difference in the methylation levels of singleton vs proximal (P-value =0.01) and a weakly significant difference between singleton vs tandem (P-value= 0.05) and singleton vs dispersed (P-value= 0.02) gene classes. No significant difference in CG methylation levels were observed between the remaining gene categories in Pigeonpea (Fig. 6E. In Soybean, we found that singleton genes showed significantly highest methylation level in comparison to all other gene categories (P-value range = 0.002 to 2.2e-16). However, we again observed that WGD genes showed lowest CG methylation levels in comparison singleton genes (P-value< 2.2e-16) and to other duplicated genes (proximal (P-value <2.2e-16; dispersed (P-value <2.2e-16)) except for tandem where we found no significant difference in methylation levels (tandem vs WGD, P-value = 0.07) (Fig. 6D).

Some studies have indicated a positive correlation between CG methylation and gene expression divergence in Cassava (H. Wang et al., 2015). However, in contrast, another study in tomato (L. Wang et al., 2018) showed a non-significant relationship between CG methylation and expression divergence. Therefore, in order to further investigate this phenomenon in Pigeonpea and Soybean, we classified duplicated gene pairs as either highly or lowly expressed, based on the fold change in the expression between gene pairs. We observed a positive association between CG body methylation and expression divergence where highly expressed gene copies have significantly (P-value = 0.02; Pigeonpea, P-value < 2.2e-16; Soybean, Wilcoxon rank sum test) high CG body methylation as compared to lowly expressed genes in both species (Fig. 6F, I). Intriguingly, CHG and CHH body methylation showed a negative correlation with expression divergence in Soybean (Fig. 6J, K). The highly expressed genes showed significantly (P-value < 0.03; CHG, P-value < 0.0001; CHH, Wilcoxon rank sum test) low CHG and CHH methylation. However, unlike Soybean, no significant difference in CHG and CHH methylation was found across the body regions of differentially expressed genes in Pigeonpea (Fig. 6G, H). Since our analysis above reveals a positive impact of CG body methylation on gene transcription in duplicated gene pairs, we further examined whether differential methylation level could contribute to expression divergence among duplicate gene pairs. To test this hypothesis, we calculated signed relative methylation (M1-M2)/(M1+M2) and expression divergence (E1-E1)/(E1+E2) in Pigeonpea and Soybean. We found a consistent positive correlation (0.07; Pigeonpea (P-value=0.006), 0.1; Soybean (P-value < 2.2e-16)) of methylation and expression divergence implying that the hypermethylated gene copy exhibited higher expression while hypomethylated copy tends to show lower expression in both species (Supporting Information Fig. S5). Overall, this significance covariation of methylation and expression divergence suggests potential role of epigenetic difference in contributing to the retention of paralog, which may further substantiate to their neofunctionalization during evolution. Moreover, we further examined whether DNA methylation influence evolution of duplicate gene pairs. To this end, average DNA methylation of paralogs genes were calculated and then correlation analysis was performed between methylation level and synonymous mutation (Ks). We found a significant negative correlation in both species (r= −0.25; Pigeonpea and r= −0.21; soybean, P-value < 2.2e-16) implicating hypermethylation in younger duplicates (Supporting Information Fig. S6).

### Natural epigenetic divergence among Pigeonpea parental and F1 hybrid line

Naturally occurring epigenetic modifications like DNA methylation remains one of the fundamental sources of heritable variations within the individuals of a species. By modulating gene expression, the stable inheritance of DNA methylation also underpins phenotypic variations often observed between genotypes(Becker et al., 2011; Raza et al., 2017; Schmitz et al., 2011; Zilberman, Gehring, Tran, Ballinger, & Henikoff, 2007). Therefore, we investigated the functional consequences of natural variation in DNA methylation in Pigeonpea. We compared the weightage methylation levels and calculated the number of DMRs in the three Pigeonpea genotypes (Fig. 7A-C). We found that, CHH DMRs were most enriched in all the three-comparison scenarios (Fig 7C). Further classification of DMRs into hyper and hypo methylated categories revealed a high abundance of hyper-methylated DMRs in F1-hybrid line relative to parental lines in all the three mC contexts (Fig. 7 A-C). However, the whole genome CG and CHG weightage methylation levels of the hybrid line were relatively low when compared to parental line. These observations are consistent with previously reported occurrence of non-additive methylation events in plants (Greaves et al., 2012; Junaid et al., 2018) where methylation levels of hybrid at specific loci is different to parents (either increased or decreased).

**Figure 7.**
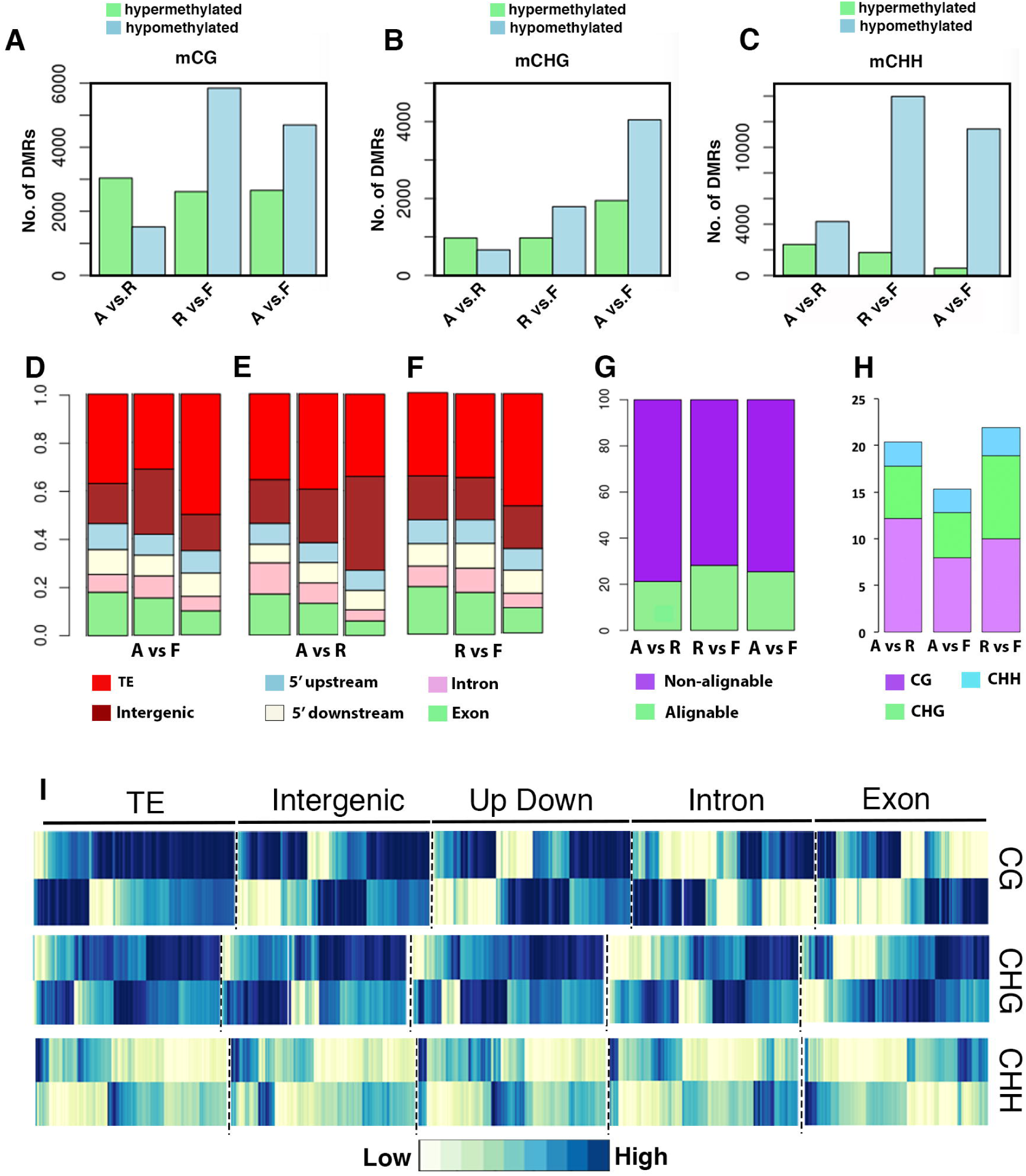
Characterization of DNA methylation variation among Pigeonpea 11A, 303R and hybrid lines: (A-C) Number of differentially methylated regions (DMRs) in (A) CG, (B) CHG and (C) CHH contexts in 11A v 303R, 11Avs Hybrid and 303R vs hybrid. (D-F). Proportions of DMRs overlapping with annotated features of genome. (G) Stacked bar plot showing percentage of DMRs in aligned vs non aligned genomic regions of Pigeon pea and Soybean. (H) Stacked bar plot showing percentage of Cg, CHG and CHH DMRs in the aligned genomic regions of Pigeon pea and Soybean. (I) Hierarchical clustering (Pearson correlation method) of methylation 11Avs303R CG, CHG and CHH DMRs in different genomic features.

We next examined the distribution of CG, CHG and CHH DMRs within the different genomic elements. We found that most of the mCG DMRs were localized in transposable element (35% in 11Avs303R; 37% in 11A vs Hybrid and 34.5% in 303R vs Hybrid) followed by gene body region (18.4% in 11Avs303R; 18% in 11A vs Hybrid; 19.7% in 303R vs Hybrid) and inter genic regions (17.8%in 11Avs303R; 16.6% in 11A vs Hybrid; 18.1% in 303R vs Hybrid) (Fig. 7D-F). A similar trend was observed for mCHG and mCHH DMRs distribution except in 303R vs hybrid where enrichment of CHH DMRs was found to be very frequent in inter genic region as compared to the rest of genomic elements. Furthermore, a subsequent hierarchical clustering of DNA methylation of DMRs revealed that TE related DMRs displayed consistently high levels of CG and CHG methylation whereas DMRs localized in the exon and intron regions showed relatively low levels of methylation (Fig. 7I). Consistent with our predictions, DMRs in CHH context showed low methylation level among all clusters.

Lastly, we investigated if the differentially methylated regions could have possibly targeted the evolutionary conserved sequences. For this, the Soybean and Pigeonpea genomes were aligned in 1:1 relationship and approximately 90 Mb sequences were found to be conserved. We found that a very low percentage of 20-28% DMRs were located in the aligned portion of genome (Fig. 7G). We next investigated the, conservation of CG, CHG and CHH DMRs in the orthologs gene and again found a very low proportion (9.8 % CG; 4.2% C-DMR) of DMRs overlapping with the 986 and 711 orthologous genes respectively (Fig. 7H). These results indicate that perhaps the variably methylated marks in the genome are not robust and thus are less likely to be conserved during evolution.

## Discussion

In-order to understand how the evolution of methylomes correlate with evolution of plant species, comparative approaches are required. In this study we performed an extensive comparative analysis of the methylomes of two closely related legume species, Pigeon pea and Soybean that diverged 20-30 million years ago, from an evolutionary perspective.

Our analysis of whole genome methylation patterns in Pigeonpea revealed higher stability of methylation in symmetric CG and CHG sites relative to asymmetric CHH sites. We surprisingly found unexpected differences in the patterns of CHH methylation in the gene and transposon density regions between the two species. Contrary to the observed general trends (Feng & Jacobsen, 2011) and to those that we observed in Soybean, CHH methylation was found to be negatively correlated in transposons rich regions and showed a positive correlation in the gene density regions of Pigeonpea genome. Such unpredicted dissimilarities in CHH methylation patterns were also found between *Brachypodium distachyon* and rice (Takuno & Gaut, 2013), two members of Poaceae family that diverged around 40-53 million years ago (International Brachypodium, 2010). These findings collectively indicate that the molecular pathways regulating CHH methylation might not always be conserved between species and may differ in some aspects.

Gene duplication is an important mechanism that leads to the generation of new genes within the genome thereby contributing novelty by paving way for neofunctionalization and evolution. However, duplicated genes have been proposed to be subjected to the constraints of “loss and gain dilemma” (Wagner, 1998, 2001) i.e., they may diverge to acquire a new function or demote to pseudogenes and ultimately face gene loss. It has also been proposed that post-duplication reduction in the expression of genes mediated by epigenetic changes contributes to a higher probability of their retention, thereby shifting the balance between loss and gain towards divergence and neofunctionalization (Chang & Liao, 2012; Rodin & Riggs, 2003). Since gene body methylation contributes to regulating expression levels, our results that gene body regions of young duplicated genes were hypermethylated and those of older duplicated genes were hypomethylated (Fig. S6) are in line with “expression divergence model”. Our findings reasonably hint towards the potential role of gene body methylation to help facilitate the retention of duplicated genes by expression regulation, in addition to transcription regulation by promoter methylations (Keller & Yi, 2014).

Gene body methylation (GbM) is an evolutionary conserved mechanism that was first discovered in crucifers (Tran et al., 2005). Although the functional roles of GbM have at large remained enigmatic, it has been proposed to play a role in regulating splicing and gene expression (Regulski et al., 2013; Tran et al., 2005; Zilberman et al., 2008).

Our analysis in Pigeonpea shows that genes with highest level of GbM showed lowest expression levels and genes with moderate GbM levels showed highest expression. These observations seem to be in agreement with a previously proposed trade off where methylation within the gene body can may curb the events of transcription initiation from cryptic promoter sites at the expense of transcriptional elongation (Zilberman et al., 2007). However, our comparative results also reveal that the association of gene expression and methylation has evolved in a complex manner during speciation, where in there is an occurrence of dosage dependent repressive role of methylation to regulate gene expression in one species, which somehow got dampened after reaching a certain threshold level by the evolutionary forces in the other species. GbM genes are characterized by some specific features such as higher gene length and slow evolutionary rates. We found that these features were maintained in both Pigeonpea and Soybean during evolution and speciation. However, our data show that Pigeonpea GbM genes tend to be slightly longer than the Soybean GbM genes. Since we also found that number of GbM genes in Soybean was higher than Pigeonpea (Fig. 3B), one possible causality of reduced gene length in Soybean may be attributed to a recent whole genome duplication event independent of Pigeonpea where, evolutionary forces may have targeted GbM gene length to balance the effect of an additional whole genome duplication on gene expression levels and to curb any possible pleiotropic effects arising due to this event in Soybean.

Examination of functional Gene ontology categories in GbM orthogous genes of Pigeon pea and showed a significant enrichment of genes involved in nitrogen compound synthesis, transport and metabolic processes as well for carbohydrate derivatives transport. These results are compelling because 1) Both Pigeon pea and Soybean are extremely important sources of protein for human nutrition and the above physiological processes are crucial in establishing protein content in pigeonpea and Soybean seeds. 2) These findings hint towards a possibility where genes involved in nitrogen and carbohydrate metabolism were repeatedly deployed or targeted for gene body methylation during evolution, to help balance their expression for optimal function. In future it will be interesting to identify specific genes in these processes and generate epi-alleles by targeted epigenome editing using CRISPR-technology (Gallego-Bartolome et al., 2018) to generate Pigeonpea and soybean varieties with higher protein content, without altering the DNA sequence.

## Acknowledgements

We thank Council of Scientific and Industrial Research, Government of India for the award of a senior research fellowship, ICAR-IARI, ICAR-NIPB and NIPB in house project, New Delhi for supporting A.J. and providing other funds and facilities.

## Author Contributions

A.J, N.K.S and K.G conceived and designed the study. A.J performed the experiments, data analysis and prepared the figures. A.J and K.G wrote the manuscript.

## References

Akalin, A., Kormaksson, M., Li, S., Garrett-Bakelman, F. E., Figueroa, M. E., Melnick, A., & Mason, C. E. (2012). methylKit: a comprehensive R package for the anal ysis of genome-wide DNA methylation profiles. Genome Biol, 13(10), R87. doi: 10.1186/gb-2012-13-10-r87

Becker, C., Hagmann, J., Muller, J., Koenig, D., Stegle, O., Borgwardt, K., & Weigel, D. (2011). Spontaneous epigenetic variation in the Arabidopsis thaliana methylome. Nature, 480(7376), 245–249. doi: 10.1038/nature10555

Bird, A. P. (1980). DNA methylation and the frequency of CpG in animal DNA. Nucleic Acids Res, 8(7), 1499–1504. doi: 10.1093/nar/8.7.1499

Carels, N., & Bernardi, G. (2000). Two classes of genes in plants. Genetics, 154(4), 1819–1825.

Chang, A. Y., & Liao, B. Y. (2012). DNA methylation rebalances gene dosage after mammalian gene duplications. Mol Biol Evol, 29(1), 133–144. doi: 10.1093/molbev/msr174

Coghlan, A., Eichler, E. E., Oliver, S. G., Paterson, A. H., & Stein, L. (2005). Chromosome evolution in eukaryotes: a multi-kingdom perspective. Trends Genet, 21(12), 673–682. doi: 10.1016/j.tig.2005.09.009

Cokus, S. J., Feng, S., Zhang, X., Chen, Z., Merriman, B., Haudenschild, C. D., … Jacobsen, S. E. (2008). Shotgun bisulphite sequencing of the Arabidopsis genome reveals DNA methylation patterning. Nature, 452(7184), 215–219. doi: 10.1038/nature06745

Domcke, S., Bardet, A. F., Adrian Ginno, P., Hartl, D., Burger, L., & Schubeler, D. (2015). Competition between DNA methylation and transcription factors determines binding of NRF1. Nature, 528(7583), 575–579. doi: 10.1038/nature16462

Drouin, G., & Dover, G. A. (1990). Independent gene evolution in the potato actin gene family demonstrated by phylogenetic procedures for resolving gene conversions and the phylogeny of angiosperm actin genes. J Mol Evol, 31(2), 132–150. doi: 10.1007/bf02109482

Du, Z., Zhou, X., Ling, Y., Zhang, Z., & Su, Z. (2010). agriGO: a GO analysis toolkit for the agricultural community. Nucleic Acids Res, 38(Web Server issue), W64–70. doi: 10.1093/nar/gkq310

Feng, S., & Jacobsen, S. E. (2011). Epigenetic modifications in plants: an evolutionary perspective. Curr Opin Plant Biol, 14(2), 179–186. doi: 10.1016/j.pbi.2010.12.002

Fultz, D., Choudury, S. G., & Slotkin, R. K. (2015). Silencing of active transposable elements in plants. Curr Opin Plant Biol, 27, 67–76. doi: 10.1016/j.pbi.2015.05.027

Gallego-Bartolome, J., Gardiner, J., Liu, W., Papikian, A., Ghoshal, B., Kuo, H. Y., … Jacobsen, S. E. (2018). Targeted DNA demethylation of the Arabidopsis genome using the human TET1 catalytic domain. Proc Natl Acad Sci U S A, 115(9), E2125–E2134. doi: 10.1073/pnas.1716945115

Greaves, I. K., Groszmann, M., Ying, H., Taylor, J. M., Peacock, W. J., & Dennis, E. S. (2012). Trans chromosomal methylation in Arabidopsis hybrids. Proc Natl Acad Sci U S A, 109(9), 3570–3575. doi: 10.1073/pnas.1201043109

Hollister, J. D., & Gaut, B. S. (2009). Epigenetic silencing of transposable elements: a trade-off between reduced transposition and deleterious effects on neighbo ring gene expression. Genome Res, 19(8), 1419–1428. doi: 10.1101/gr.091678.109

Hollister, J. D., Smith, L. M., Guo, Y. L., Ott, F., Weigel, D., & Gaut, B. S. (2011). Transposable elements and small RNAs contribute to gene expression divergence between Arabidopsis thaliana and Arabidopsis lyrata. Proc Natl Acad Sci U S A, 108(6), 2322–2327. doi: 10.1073/pnas.1018222108

International Brachypodium, I. (2010). Genome sequencing and analysis of the model grass Brachypodium distachyon. Nature, 463(7282), 763–768. doi: 10.1038/nature08747

Jiang, N., Bao, Z., Zhang, X., Eddy, S. R., & Wessler, S. R. (2004). Pack-MULE transposable elements mediate gene evolution in plants. Nature, 431(7008), 569–573. doi: 10.1038/nature02953

Junaid, A., Kumar, H., Rao, A. R., Patil, A. N., Singh, N. K., & Gaikwad, K. (2018). Unravelling the epigenomic interactions between parental inbreds resulting in an altered hybrid methylome in pigeonpea. DNA Res, 25(4), 361–373. doi: 10.1093/dnares/dsy008

Keller, T. E., & Yi, S. V. (2014). DNA methylation and evolution of duplicate genes. Proc Natl Acad Sci U S A, 111(16), 5932–5937. doi: 10.1073/pnas.1321420111

Kim, D., Pertea, G., Trapnell, C., Pimentel, H., Kelley, R., & Salzberg, S. L. (2013). TopHat2: accurate alignment of transcriptomes in the presence of insertions, deletio ns and gene fusions. Genome Biol, 14(4), R36. doi: 10.1186/gb-2013-14-4-r36

Kim, K. D., El Baidouri, M., Abernathy, B., Iwata-Otsubo, A., Chavarro, C., Gonzales, M., … Jackson, S. A. (2015). A Comparative Epigenomic Analysis of Polyploidy-Derived Genes in Soybean and Common Bean. Plant Physiol, 168(4), 1433–1447. doi: 10.1104/pp.15.00408

Kim, K. D., Shin, J. H., Van, K., Kim, D. H., & Lee, S. H. (2009). Dynamic rearrangements determine genome organization and useful traits in soybean. Plant Physiol, 151(3), 1066–1076. doi: 10.1104/pp.109.141739

Lang, Z., Wang, Y., Tang, K., Tang, D., Datsenka, T., Cheng, J., … Zhu, J. K. (2017). Critical roles of DNA demethylation in the activation of ripening-induced genes and inhibition of ripening-repressed genes in tomato fruit. Proc Natl Acad Sci U S A, 114(22), E4511–E4519. doi: 10.1073/pnas.1705233114

Lavin, M., Herendeen, P. S., & Wojciechowski, M. F. (2005). Evolutionary rates analysis of Leguminosae implicates a rapid diversification of lineages during the tertiary. Syst Biol, 54(4), 575–594. doi: 10.1080/10635150590947131

Lister, R., O’Malley, R. C., Tonti-Filippini, J., Gregory, B. D., Berry, C. C., Millar, A. H., & Ecker, J. R. (2008). Highly integrated single-base resolution maps of the epigenom e in Arabidopsis. Cell, 133(3), 523–536. doi: 10.1016/j.cell.2008.03.029

Liu, R., How-Kit, A., Stammitti, L., Teyssier, E., Rolin, D., Mortain-Bertrand, A., … Gallusci, P. (2015). A DEMETER-like DNA demethylase governs tomato fruit ripening. Proc Natl Acad Sci U S A, 112(34), 10804–10809. doi: 10.1073/pnas.1503362112

Nei, M., & Gojobori, T. (1986). Simple methods for estimating the numbers of synonymous and nonsynonymous nucleotide substitutions. Molecular Biology and Evolution, 3(5), 418–426. doi: 10.1093/oxfordjournals.molbev.a040410

Niederhuth, C. E., Bewick, A. J., Ji, L., Alabady, M. S., Kim, K. D., Li, Q., … Schmitz, R. J. (2016). Widespread natural variation of DNA methylation within angiosperms. Genome Biol, 17(1), 194. doi: 10.1186/s13059-016-1059-0

Nuthikattu, S., McCue, A. D., Panda, K., Fultz, D., DeFraia, C., Thomas, E. N., & Slotkin, R. K. (2013). The initiation of epigenetic silencing of active transposable elements is triggered by RDR6 and 21-22 nucleotide small interfering RNAs. Plant Physiol, 162(1), 116–131. doi: 10.1104/pp.113.216481

Otto, S. P., & Whitton, J. (2000). Polyploid incidence and evolution. Annu Rev Genet, 34, 401–437. doi: 10.1146/annurev.genet.34.1.401

Paterson, A. H., Freeling, M., Tang, H., & Wang, X. (2010). Insights from the comparison of plant genome sequences. Annu Rev Plant Biol, 61, 349–372. doi: 10.1146/annurev-arplant-042809-112235

Pfeifer, G. P. (2006). Mutagenesis at methylated CpG sequences. Curr Top Microbiol Immunol, 301, 259–281. doi: 10.1007/3-540-31390-7_10

Raza, M. A., Yu, N., Wang, D., Cao, L., Gan, S., & Chen, L. (2017). Differentia l DNA methylation and gene expression in reciprocal hybrids between Solanum lycopersicum and S. pimpinellifolium. DNA Research, 24(6), 597–607. doi: 10.1093/dnares/dsx028

Regulski, M., Lu, Z., Kendall, J., Donoghue, M. T., Reinders, J., Llaca, V., … Martienssen, R. A. (2013). The maize methylome influences mRNA splice sites and reveals widespread paramutation-like switches guided by small RNA. Genome Res, 23(10), 1651–1662. doi: 10.1101/gr.153510.112

Robertson, K. D. (2005). DNA methylation and human disease. Nat Rev Genet, 6(8), 597–610. doi: 10.1038/nrg1655

Rodin, S. N., & Riggs, A. D. (2003). Epigenetic silencing may aid evolution by gene duplication. J Mol Evol, 56(6), 718–729. doi: 10.1007/s00239-002-2446-6

Schmitz, R. J., Schultz, M. D., Lewsey, M. G., O’Malley, R. C., Urich, M. A., Libiger, O., … Ecker, J. R. (2011). Transgenerational epigenetic instability is a sour ce of novel methylation variants. Science, 334(6054), 369–373. doi: 10.1126/science.1212959

Schmutz, J., Cannon, S. B., Schlueter, J., Ma, J., Mitros, T., Nelson, W., … Jacks on, S. A. (2010). Genome sequence of the palaeopolyploid soybean. Nature, 463(7278), 178–183. doi: 10.1038/nature08670

Serres-Giardi, L., Belkhir, K., David, J., & Glemin, S. (2012). Patterns and evolution of nucleotide landscapes in seed plants. Plant Cell, 24(4), 1379–1397. doi: 10.1105/tpc.111.093674

Seymour, D. K., & Gaut, B. S. (2019). Phylogenetic shifts in gene body methylation correlate with gene expression and reflect trait conservation. Mol Biol Evol. doi: 10.1093/molbev/msz195

Slotkin, R. K., & Martienssen, R. (2007). Transposable elements and the epi genetic regulation of the genome. Nat Rev Genet, 8(4), 272–285. doi: 10.1038/nrg2072

Takuno, S., & Gaut, B. S. (2012). Body-methylated genes in Arabidopsis thaliana are functionally important and evolve slowly. Mol Biol Evol, 29 (1), 219–227. doi: 10.1093/molbev/msr188

Takuno, S., & Gaut, B. S. (2013). Gene body methylation is conserved between plant orthologs and is of evolutionary consequence. Proc Natl Acad Sci U S A, 110(5), 1797–1802. doi: 10.1073/pnas.1215380110

Takuno, S., Seymour, D. K., & Gaut, B. S. (2017). The Evolutionary Dynamics of Orthologs That Shift in Gene Body Methylation between Arabidopsis Species. Mol Biol Evol, 34(6), 1479–1491. doi: 10.1093/molbev/msx099

Thompson, J. D., Higgins, D. G., & Gibson, T. J. (1994). CLUSTAL W: improving the sensitivity of progressive multiple sequence alignment through sequence weighting, position-specific gap penalties and weight matrix choice. Nucleic Acids Res, 22(22), 4673–4680. doi: 10.1093/nar/22.22.4673

Tran, R. K., Henikoff, J. G., Zilberman, D., Ditt, R. F., Jacobsen, S. E., & Henikoff, S. (2005). DNA methylation profiling identifies CG methylation clusters in Arabidopsis genes. Curr Biol, 15(2), 154–159. doi: 10.1016/j.cub.2005.01.008

Varshney, R. K., Chen, W., Li, Y., Bharti, A. K., Saxena, R. K., Schlueter, J. A., … Jackson, S. A. (2011). Draft genome sequence of pigeonpea (Cajanus cajan), an orphan legume crop of resource-poor farmers. Nat Biotechnol, 30(1), 83–89. doi: 10.1038/nbt.2022

Wagner, A. (1998). The fate of duplicated genes: loss or new function? Bioessays, 20(10), 785–788. doi: 10.1002/(SICI)1521-1878(199810)20:10<785::AID-BIES2>3.0.CO;2-M

Wagner, A. (2001). Birth and death of duplicated genes in completely sequenced eukaryotes. Trends Genet, 17(5), 237–239. doi: 10.1016/s0168-9525(01)02243-0

Wang, H., Beyene, G., Zhai, J., Feng, S., Fahlgren, N., Taylor, N. J., … Ausin, I. (2015). CG gene body DNA methylation changes and evolution of duplicated genes in cassava. Proc Natl Acad Sci U S A, 112(44), 13729–13734. doi: 10.1073/pnas.1519067112

Wang, L., Xie, J., Hu, J., Lan, B., You, C., Li, F., … Wang, H. (2018). Comparative epigenomics reveals evolution of duplicated genes in potato and tomato. Plant J, 93(3), 460–471. doi: 10.1111/tpj.13790

Wang, Y., Tang, H., DeBarry, J. D., Tan, X., Li, J., Wang, X., … Paterson, A. H. (2012). MCScanX: a toolkit for detection and evolutionary analysis of gene synteny and collinearity. Nucleic Acids Research, 40(7), e49–e49. doi: 10.1093/nar/gkr1293

Wolfe, K. H. (2001). Yesterday’s polyploids and the mystery of diploidization. Nat Rev Genet, 2(5), 333–341. doi: 10.1038/35072009

Xi, Y., & Li, W. (2009). BSMAP: whole genome bisulfite sequence MAPping program. BMC Bioinformatics, 10, 232. doi: 10.1186/1471-2105-10-232

Yang, L., Takuno, S., Waters, E. R., & Gaut, B. S. (2011). Lowly expressed genes in Arabidopsis thaliana bear the signature of possible pseudogenization by promoter degradation. Mol Biol Evol, 28(3), 1193–1203. doi: 10.1093/molbev/msq298

Zhang, X., Yazaki, J., Sundaresan, A., Cokus, S., Chan, S. W., Chen, H., … Ecker, J. R. (2006). Genome-wide high-resolution mapping and functional analysis of DNA methylation in arabidopsis. Cell, 126(6), 1189–1201. doi: 10.1016/j.cell.2006.08.003

Zhang, Z., Li, J., Zhao, X.-Q., Wang, J., Wong, G. K.-S., & Yu, J. (2006). KaKs_Calculator: Calculating Ka and Ks Through Model Selection and Model Averaging. Genomics, Proteomics & Bioinformatics, 4(4), 259–263. doi: https://doi.org/10.1016/S1672-0229(07)60007-2

Zhang, Z., Xiao, J., Wu, J., Zhang, H., Liu, G., Wang, X., & Dai, L. (2012). ParaAT: A parallel tool for constructing multiple protein-coding DNA alignments. Biochemica l and Biophysical Research Communications, 419(4), 779–781. doi: https://doi.org/10.1016/j.bbrc.2012.02.101

Zhu, H., Wang, G., & Qian, J. (2016). Transcription factors as readers and effectors of DNA methylation. Nat Rev Genet, 17(9), 551–565. doi: 10.1038/nrg.2016.83

Zilberman, D., Coleman-Derr, D., Ballinger, T., & Henikoff, S. (2008). Histone H 2A.Z and DNA methylation are mutually antagonistic chromatin marks. Nature, 456(7218), 125–129. doi: 10.1038/nature07324

Zilberman, D., Gehring, M., Tran, R. K., Ballinger, T., & Henikoff, S. (2007). Genome-wide analysis of Arabidopsis thaliana DNA methylation uncovers an interdependence between methylation and transcription. Nat Genet, 39(1), 61–69. doi: 10.1038/ng1929

